# Where do our graduates go? A toolkit for retrospective and ongoing career outcomes data collection for biomedical PhD students and postdoctoral scholars

**DOI:** 10.1101/539031

**Authors:** Elizabeth A. Silva, Alicia B. Mejía, Elizabeth S. Watkins

**Affiliations:** Graduate Division, University of California, San Francisco, San Francisco, California, United States of America

## Introduction

Universities are at long last undertaking efforts to collect and disseminate information about student career outcomes, after decades of calls to action. Organizations such as Rescuing Biomedical Research and Future of Research brought this issue to the forefront of graduate education, and the second Future of Biomedical Graduate and Postdoctoral Training conference (FOBGAPT2) featured the collection of career outcomes data in its final recommendations, published in 2017 (Hitchcock et al., 2017). More recently, 30 institutions assembled as the Coalition for Next Generation Life Science, committing to ongoing collection and dissemination of career data for both graduate and postdoc alumni. A few individual institutions have shared snapshots of the data in peer-reviewed publications (Mathur et al., 2018; Silva, Jarlais, Lindstaedt, Rotman, & Watkins, 2016) and on websites. As more and more institutions take up this call to action, they will now be looking for tools, protocols, and best practices for ongoing career outcomes data collection, management, and dissemination. Here, we describe UCSF’s experiences in conducting a retrospective study, and in institutionalizing a methodology for annual data collection and dissemination. We describe and share all tools we have developed, and we provide calculations of the time and resources required to accomplish both retrospective studies and annual updates. We also include broader recommendations for implementation at your own institutions, increasing the feasibility of this endeavor.

## Global Recommendations

### DON’T LET THE PERFECT BE THE ENEMY OF THE GOOD

Exceedingly long planning stages have kept career outcomes collection and reporting on institutional back burners for decades. We acknowledged from the outset that we would not be able to find every alumnus or to categorize every job title with precision. We chose a repository with sufficient flexibility so that post-hoc adjustments to the data would be feasible. We also made our peace with missing information in the retrospective study, knowing that the quantity and quality of data will improve as we add new graduates to the dset in the on-going collection.

### DEVELOP A PROJECT CHARTER

A project charter sets boundaries on the scope and scale of the project, articulates the roles of the personnel, and includes a timeline and description of the project milestones. This document was critical for ensuring the project progressed at an acceptable pace and for preventing “mission creep” - unplanned expansions that present barriers to completion. A charter was particularly important for our postdoc dataset. For decades there has been a dearth of data about postdocs (National Academy of Sciences, National Academy of Engineering, and Institute of Medicine. 2014). As the project grew so did the enthusiasm for expanding to include additional data-points that were not directly relevant to the objectives of the project (eg. date of birth, time in previous postdoc). While these data-points are interesting and valuable, defined limits on the scope of the project are necessary for completion. We provide our project charter in supporting information file S1 as a sample.

### IDENTIFY CAMPUS STAKEHOLDERS

On every campus, career outcomes data are collected and reported by a variety of stakeholders, often with little coordination of efforts and resources. Coordination with stakeholders offers the opportunity to improve the quality of the dataset while reducing the overall institutional resources required. *Graduate programs*: Individual graduate program staff and faculty often have first-hand knowledge of the current positions of graduates, having maintained personal connections years after graduation. Programs may or may not have developed a stable repository or platform for storing and reporting alumni information. Collaboration with the graduate programs involves collecting accurate alumni information and offering a central platform along with user support for accessing the data. *T32 program directors:* In applying for and reporting progress for the Ruth L. Kirschstein Institutional National Research Service Award, Principle Investigators are required to report first position and current position for every funded trainee for 15 years. Meeting this requirement is an enormous undertaking and is resource intensive. Equally, there is a great deal of data contained in the reports that can be extracted. *Alumni relations:* Alumni relations generously shared email contact information from alumni in their database to assist with our survey. We reported our survey response rates to alumni relations, who reported the responses as successful touchpoints.

## Data Overview

We have developed and are maintaining two distinct datasets, one for PhD alumni (Figure 1) and one for postdoctoral alumni (not shown). Our PhD alumni dataset includes every student who began a PhD program at UCSF since 1996. A record is created for the student as they matriculate to the program, rather than as they graduate. Our postdoctoral alumni dataset includes every postdoctoral scholar (postdoc) who left the institution since 2011. In both cases we include all available demographic information, previous education, program and degree information, and job titles and employers. Data is transferred from the student information system (PhD alumni, via application program interface [API]) and the Office of Institutional Research (postdoctoral alumni, based on Human Resources records). Career information is collected annually for up to 15 years after a student or postdoc leaves the institution, but is displayed on the public website in 5 year increments and/or 5 year aggregates (https://graduate.ucsf.edu/program-statistics; https://postdocs.ucsf.edu/postdocs-ucsf). A full description of all metadata for the PhD dataset is provided in supporting information file S2 and for the postdoc dataset is provided in supporting information file S3.

**Figure 1.**
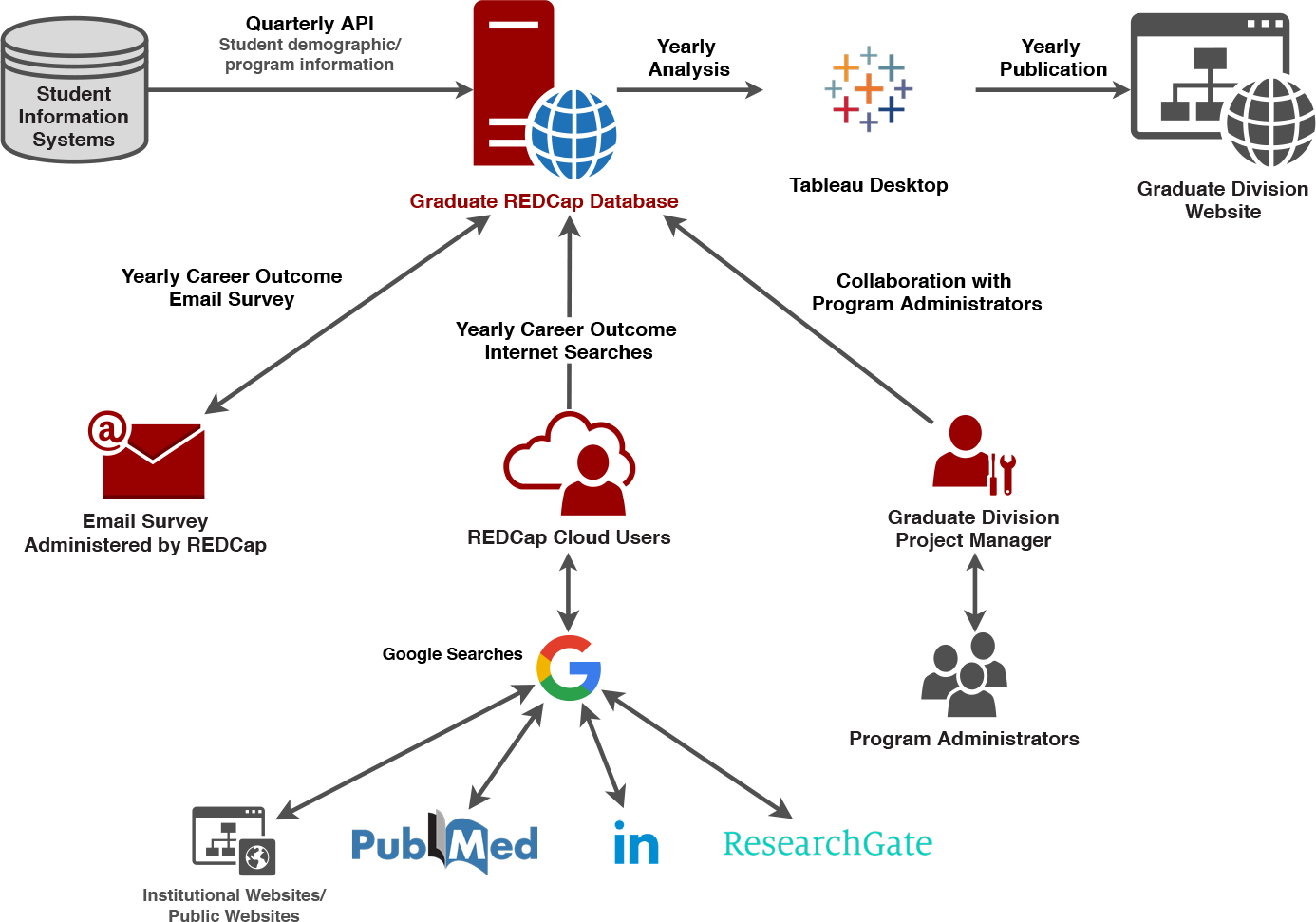
Overview of data flow for the PhD alumni career outcomes. Postdoctoral outcomes data flow (not shown) is similar except where noted below. Basic demographic and degree information is transferred to REDCap from the student information system (postdoctoral data comes from human resources). Annually, staff administer a one-time survey requesting current employment information from the PhD alumni, and conduct online searches for those who do not return a survey (postdocs and PhDs). Employment data is recorded in REDcap. Data is then uploaded to Tableau for public display on the graduate division (or postdoctoral) website. PhD data is also shared with the PhD program staff: staff provide updates to the project manager about program alumni, and data collected by our team is shared with the program staff

## Setting up a repository

We considered multiple systems and platforms for our data collection, management, curation, and archiving, including MS Excel, MS Access, Smartsheets, Salesforce, and REDCap. We considered the following features in our analysis:

Required:

- Cloud based, to allow multiple users
- Compatible with Mac and PC, to allow multiple users
- No requirement for individual user license, to enable access by multiple users
- Export and import via comma separated value (.CSV) files

Recommended:

- Variable user permissions, to allow access for stakeholders
- Flexible data fields, as far as possible
- Open source, or otherwise accessible for additional development work
- Survey option
- Report builder

Ultimately, we determined that flexibility and stability were most crucial, so we could adjust fields as we gained a better understanding of what the data were like, and so we would not risk corrupting the data. We opted for REDCap, developed by Vanderbilt University, which is a free and open source and includes all of our required features (https://www.project-redcap.org/). REDCap allows for “data access groups”, which means different users can be given different access to subsets of data as defined by the administrator, including view-only. This flexibility allows various stakeholders on campus to access the datasets relevant to them without violating the Federal Education Rights and Privacy Act (FERPA) and without risking corruption of the data. It can also send surveys (more on that below) and can generate reports that can be exported as .CSV files. As an open-source database, REDCap allows for the development of APIs for data import and export. We took advantage of this feature and updated our student demographic and enrollment data directly from UCSF’s student information system.

Here we provide our REDCap data dictionaries for our graduate student outcomes database (S4) and our postdoc outcomes database (S5). These data dictionaries can be used to re-create an empty database in REDCap, which can then be modified according to the needs and interests of your institution.

## Outcomes data collection and curation

There are two phases to the data collection effort: retrospective and ongoing. We describe each separately.

### RETROSPECTIVE DATA COLLECTION: CYBER-SLEUTHING

We began with a retrospective study in 2017, relying entirely on internet searches (cyber-sleuthing). This method was previously described (Silva et al., 2016). Since this publication, we have accrued new best practices. First, while LinkedIn is a superior platform for gathering career information, particularly for individuals in the private sector, Google is a superior search engine. A search for [First Name] [Last Name] “LinkedIn” is more likely to yield relevant results.

Additionally, Google’s search results can be influenced by logging into a LinkedIn account for a user who is well-connected to your alumni. Google’s enriched results incorporate user information (sites visited, current location, logins to social media accounts) in an effort to return the most relevant information. When a user is logged into a LinkedIn account with more connections to institutional alumni, Google is more likely to return top hits for institutional alumni. Author E. Silva has 800+ LinkedIn connections, many of who are current UCSF staff and students, or UCSF alumni. When she is logged into her account, relevant search results were more forthcoming than for team members with fewer connections. Furthermore, identification of individuals as 2nd or 3rd order connections to the LinkedIn user serves as verification and helps disambiguate individuals with similar names. We recorded one position per year for up to 15 years after leaving the institution.

### ONGOING COLLECTION: PHD ALUMNI SURVEYS AND CYBER-SLEUTHING

In 2018 we set out to update our datasets, recording the most recent position for each alumnus. For our PhD alumni dataset we introduced survey results into our data collection method. We sent a simple 4-question survey to all PhD program alumni for whom we had an email address:

- What is your job title?
- What is the name of your organization/institution/company?
- City
- State (or Country if not US)

This survey was sent to 1732 alumni with a functioning email address, and 800 responses were received, for a return of 43%. Overall this represents 30% of our alumni. We attribute the high response rate to two factors: (1) brevity of the survey and (2) an appeal to the cause. The email inviting alumni to participate stated that the survey would take less than one minute to complete and explained that the data collected would be used for transparent and thorough reporting of career outcomes. Respondents were also assured that data would be displayed only in aggregate and anonymously. We included a link to our public display of retrospective data so that prospective participants could see how the data were used. Once the survey was complete, we updated the remaining 70% of PhD alumni through cyber-sleuthing.

While we publicly display results in five-year increments, we identified three significant advantages to annual data collection. First, it is easier to find and update each individual annually via the cyber-sleuthing method. Second, annually updated data will show more-nuanced career trajectories, which will assist our student and postdoctoral services staff as they advise trainees on career exploration and decision-making. Third, the National Institutes of Health require that institutional training grants (T32) awardees provide annual updates of the career outcomes of funded trainees, for which these data can serve as a resource.

### CLASSIFICATION

We use the taxonomy developed collectively in 2017 by representatives of universities with NIH Broadening Experiences in Scientific Training (BEST) awards, members of Rescuing Biomedical Research (RBR) and the founding institutions of the Coalition for Next Generation Life Science. (CNGLS) Classification terms were applied by our staff, rather than the alumni themselves, in an effort to ensure consistency. Most positions fall clearly into categories for career type and section; however, many jobs do not fall clearly in a specific career category for job function. When a position did not clearly fall into a category, we discussed its best placement as a group and then added further notes to the definitions associated with each category to clarify how the categories should be applied (S6). Once initial classification was complete, we randomly assigned a subset of records for re-review by coders – those who applied the classifications to the alumni. Two hundred individuals were assigned to each of three reviewers. Using a basic spreadsheet, each reviewer indicated records the issue. In this process, we identified a few errors, but more importantly we identified inconsistencies in coding that could be rectified in bulk. For example, a number of institutions, including UCSF, have fellows’ programs that provide a pathway from graduate school to independent research, effectively skipping the postdoctoral stage. Our team had discrepant understandings of whether to classify these positions as training positions, or independent faculty-like positions (“faculty, tenure-track not applicable”). The audit highlighted the discrepancy and prompted classification that might require review and provided notes describing the issue. In this process, we identified a few errors, but more importantly we identified inconsistencies in coding that could be rectified in bulk. For example, a number of institutions, including UCSF, have fellows’ programs that provide a pathway from graduate school to independent research, effectively skipping the postdoctoral stage. Our team had discrepant understandings of whether to classify these positions as training positions, or independent faculty-like positions (“faculty, tenure-track not applicable”). The audit highlighted the discrepancy and prompted classification decisions. Any necessary re-classifications were then extended to the full dataset. A summary of the audit is provided in Table 1.

**Table 1.** Summary of data audit for PhD and postdoctoral alumni data.

| Post-collection Data Audit Statistics | PhD alumni | Postdoctoral alumni |
| --- | --- | --- |
| Total Trainees | 2,557 | 2,355 |
| Total Entries | 16,084 | 12,921 |
| Total trainees reviewed | 546 | 531 |
| Total entries reviewed | 3,536 | 2,576 |
| Total Trainees corrected | 49 | 191 |
| Total Number of Corrections Identified (Entries) | 153 | 538 |
| Correction type and frequency: | Tenure Track - 89<br>Other Data Entry - 20<br>Other Classification Error - 20<br>Added/Corrected Link - 19<br>Other Misc - 2<br>Trainee Information from SIS - 1<br>Group Leader - 1<br>Entrepreneur - 1 | Added/Corrected Link - 199*<br>Other Classification Error - 95<br>Tenure Track - 77<br>Other Data Entry - 51<br>Group Leader - 50<br>UCSF Associate/Assistant<br>Specialist/Researcher - 24<br>UCSF Title - 21<br>Trainee Information from OIR - 12<br>Entrepreneur - 8<br>Other Misc - 1 |
| Total Number of Corrections Accepted (Entries) | 73 | 427 |
| % Corrections Accepted to Sample Size | 2% | 17% |
| % Corrections Accepted to Population Size | 0.45% | 3% |

\*URL for LinkedIn, institutional website, or other Internet site where alumnus information is available

### UNKNOWNS

Some people cannot be reached by email or cannot be located online. For example, individuals who are unemployed rarely identify as such. We observed that those working in clinical practice are disproportionately difficult to find online since they neither use LinkedIn nor have comprehensive profile pages on institutional websites. Alumni who left the institution more recently are easier to find, and current position is easier to find than any past-held position. In Table 2 we summarize the proportion of PhD alumni for whom we were unable to find information (unknowns) in our retrospective study, comparing current position (2017) to past-held positions (1996-2016).

**Table 2.** Number and percent of PhD alumni for whom data was missing in our initial retrospective study (2017).

| Cohort | Count of Trainees | % No Job Title* 1996-2016 | Completely Unknown** 1996-2016 | % Completely Unknown** 1996-2016 | % No Job Title 2017*** | Completely Unknown 2017**** | % Completely Unknown 2017**** |
| --- | --- | --- | --- | --- | --- | --- | --- |
| 1996 | 83 | 39% | 3 | 4% | 17% | 2 | 5% |
| 1997 | 77 | 45% | 6 | 8% | 17% | 3 | 6% |
| 1998 | 102 | 35% | 14 | 14% | 3% | 3 | 3% |
| 1999 | 98 | 38% | 6 | 6% | 16% | 9 | 10% |
| 2000 | 122 | 35% | 10 | 8% | 19% | 13 | 11% |
| 2001 | 108 | 23% | 2 | 2% | 12% | 3 | 3% |
| 2002 | 142 | 28% | 12 | 8% | 16% | 22 | 15% |
| 2003 | 147 | 22% | 5 | 3% | 12% | 7 | 5% |
| 2004 | 127 | 24% | 6 | 5% | 13% | 11 | 9% |
| 2005 | 153 | 15% | 0 | 0% | 10% | 2 | 1% |
| 2006 | 128 | 16% | 11 | 9% | 10% | 13 | 10% |
| 2007 | 137 | 16% | 1 | 1% | 9% | 1 | 1% |
| 2008 | 116 | 10% | 1 | 1% | 9% | 4 | 3% |
| 2009 | 96 | 13% | 0 | 0% | 11% | 0 | 0% |
| 2010 | 70 | 13% | 2 | 3% | 10% | 2 | 3% |
| 2011 | 50 | 18% | 3 | 6% | 12% | 3 | 6% |
| 2012 | 38 | 18% | 3 | 8% | 19% | 4 | 11% |
| Total | 1794 | 24% | 85 | 5% | 12% | 102 | 6% |

\*Job title not found for previous years \*\*No information found for previous years (job title, organization, location) \*\*\*Job title not found for 2017 (current position at the time of search) \*\*\*\*No information found for 2017 (current job title, organization, location at the time of search)

## Resources needed

The scope and scale of this project demanded significant staff time. Here, we estimate the amount of time required and describe the roles and responsibilities of the primary personnel. We also provide a more detailed summary of our timeline, milestones, and team members in our charter document (below and S6).

### PRIMARY PERSONNEL

#### Project sponsor

Decision maker for the overall project, directs data collection and analysis.

#### Project/data manager

Documents project goals, documents and communicates project status, tracks time and effort spent, identifies roles and responsibilities, and monitors other project details. Secondary roles include data collection, consolidation and management in REDCap, database administration, and data quality audits and cleanup.

#### Project support

Undergraduate student intern. Collects and consolidates career outcomes in REDCap, and classifies the job titles and employers.

### TIMELINE AND MILESTONES

The data collection and classification for our 15-year retrospective study of PhD student alumni, undertaken in 2017, was completed in three months (June 15 to September 15). Through the remainder of 2017 and into 2018, a project sustainment plan was developed and implemented by the project manager, and the project was expanded to include a retrospective study of the postdoc population. An update of all PhD and postdoc alumni outcomes was completed in the three summer months of 2018. In supporting file S7, we provide a worksheet that estimates the resources that would be required at other institutions for a retroactive data search and for an annual update of the alumni outcomes.

## Summary

Many institutions report that they have delayed commitment to these projects due to concerns about the resources required. Having done the work to implement systems for retrospective and on-going data collection, we share all of our materials and resources here to motivate other institutions to take up this call to action. Transparency in career outcomes for PhD students and graduates is an achievable goal and, we argue, a responsibility that universities must fulfill.

