## Supplemental file 1 for "Where do our graduates go? A toolkit for retrospective and ongoing career outcomes data collection for biomedical PhD students and postdoctoral scholars"

### PhD Career Outcomes Project Charter

UCSF GRADUATE DIVISION

Created by UCSF Graduate Division

### Historical Career Outcomes Collection Project

#### Project Background

There is a great need for better assessment of career outcomes for graduate students and postdoctoral scholars due to the oversupply of postdoctoral scholars. A study of postdoctoral outcomes at the University of California, San Francisco suggests that institutions have an obligation to determine where their postdoc alumni are employed and to share this information with current and future trainees. Currently the Ruth L. Kirschstein National Research Service Award (T32) funding scheme of the National Institutes of Health require that an academic institution report the current employment status and positions for all students and postdocs each year, with required trainee follow-up for up to 15 years after leaving UCSF. This information, however is not made readily available to graduate and post-doc students in a form to weigh their options in planning and preparing for their future careers.

This study has proposed that institutions must accept responsibility for providing better transparency of career outcomes to the fullest extent possible at a local level. With a more robust collection and analysis career outcomes, institutions can easily provide an abundance of existing and freely available information, resources, strategies, and curricula. This information can serve immediately as a career development resource, allowing students to become aware of the opportunities that are available to them. This project has been initiated to pilot the more comprehensive collection, consolidation and analysis of the career and demographic information of UCSF PhD students for a period of 15 years post-graduation, for the purpose of understanding and communicating the career trajectories of UCSF PhD students.

Created by UCSF Graduate Division

#### Problem Statement/ Business Needs

- Each UCSF graduate program provides a separate report on the current employment information for all students and postdocs for each year. This creates unnecessary variability in reporting style and redundancy in work effort.
- More comprehensive demographic and job status/position information should be collected and analyzed regarding the career trajectories of UCSF graduates for the purpose of guiding and informing current students of their career options.
- Career trajectory statistics are not made available to the UCSF community or the general public.
- A retrospective study should be initiated, as it may take several years to create a sample size large enough to produce reliable trends due to the small yearly graduate student populations.
- There is no current model upon which we can rely for collecting and disseminating outcomes, so this initiative will need to serve as a guide for institutes nationwide.

#### Objectives

- Collect compulsory reporting information from all the graduate program offices, where feasible, and amalgamate into a single database repository. This will eliminate redundant reporting activities and standardize the content and format of the information provided to the NIH by the programs.

- Associate more comprehensive demographic data describing each graduate student.
- Collect current employment status and position, as well as retrospective information for up to 15 years after a graduate student has graduated from UCSF.
- Categorize resultant data collected regarding employment and job position into appropriate sector, job type, and job function.
- Analyze the present career trajectory outcomes and statistics on the Graduate Division website.
- Develop and document a Sustainment Plan for ongoing collection, consolidation and analysis of student career information. Student career outcomes will be provided to the T32 Trainee Tracking Manager, Halima Mohammed, who will report to the NIH.
- Document a process or best practice for other institutions to follow.

### Scope

#### In-Scope

1. Collection of demographic data and career outcomes for UCSF PhD students who have matriculated since 1996 to Present, from the following basic science and social & population science programs. Yearly career outcomes are collected for up to 15 years after graduation. Some students do not complete the PhD program, but are included in this scope as having started with the intention of completing a PhD program.
    1. Tetrad (Biochemistry and Molecular Biology, Cell Biology, Genetics).
    2. Bioengineering
    3. Biological and Medical Informatics
    4. Biomedical Sciences
    5. Biophysics
    6. Chemistry and Chemical Biology
    7. Developmental and Stem Cell Biology
    8. Neuroscience
    9. Oral and Craniofacial Sciences
    10. Pharmaceutical Sciences and Pharmacogenomics
    11. Epidemiology and Translational Science
    12. Global Health Sciences
    13. History of Health Sciences
    14. Medical Anthropology
    15. Nursing
    16. Sociology
2. Publication of PhD demographic and career outcome findings on Graduate Division website with links to OIR website.
3. Create a database to facilitate the collection, storage, reporting and analysis of the demographic and career profiles of UCSF PhD graduate students.

#### Out-of-Scope

1. Students of the Professional Doctorate (Doctor of Nursing Practice and Doctor of Physical Therapy) and Certificate Programs.
2. UCSF postdoctoral scholars
3. Students in the Master’s programs
4. Anything not specifically described and included in “In Scope.”

Created by UCSF Graduate Division

#### Timeline & Milestones

| Phase | Time Frame |
|---|---|
| **Planning**<br>• Draft and approve project charter and/or project plan/timeline.<br>• Design and create Career Outcomes REDCap database/ or data collection medium<br>• Determine and document the flow of data and information | 1 month |
| **Data Consolidation & Collection**<br>• Complete upload/organization of historical trainee demographic and program/appointment information<br>• Collection of yearly career outcomes for trainees<br>○ All Cohorts determined In Scope.<br>○ Each year since graduation/separation to present year (or at least 1, 5, 10 and 15 year(s) out) | 3 months (timeline depends on number of cohorts) |
| **Data Management, Storage & Communication**<br>• Investigate data management for ongoing collection, storage, and reporting of career outcomes (if not using REDCap or pre-determined tool<br>• Establish API Import of quarterly/monthly reports for new and graduating/separating trainees | 2 months (timeline depends on current data management solutions utilized for trainees) |
| **Data Analysis & Publication**<br>• Design and publication of demographic and career outcome data in Tableau | 2 months (separate timelines for demographics and career outcomes for both PhD and Postdoc dictated by NGLS Coalition) |

Created by UCSF Graduate Division

#### Risk Summary

| Risk | Mitigation/ Contingency |
|---|---|
| Insufficient project resources to complete data collection and consolidation | Mitigation/Reduction Strategy - Project Manager to create a log/audit of all expected spreadsheets for each of the T32, including the number of students represented. This log will note which have or have not been received. |
| Data entry and copy/paste of student information into spreadsheet may contain human errors that affect the accuracy and reliability of the results of this project | Mitigation/Reduction Strategy -<br>- Project Manager to create automatic checks that data is being entered in completely and accurately<br>- Project Manager to task the Project Sponsor and SME to review and audit of sample data collected and classified for accuracy |
| Receiving complete dataset(s) from SIS to create and complete the dataase of students for collecting outcomes may be delayed. May not have a complete list of trainees on which to collect outcomes, and cannot distinguish between students currently working on a degree | Mitigation/Reduction Strategy –<br>- Project Sponsor and Project Manager to collaborate more closely with Applications Programmers at Student Information Services (Registrar) to design more reliable reports for API upload into REDCap database. |

| Risk | Mitigation/ Contingency |
|---|---|
| and those that withdrew from a program without degrees. | |

### Assumptions

- Historical career outcome information will be most efficiently collected via internet searches. Career outcome data is also available from Program Administrators, as current and first job placements are reported for T32 grant administration.
- Accuracy of collection and classification is entrusted to the Student Intern/Project Support.

### Dependencies

- Career taxonomy is finalized and communicated by the Coalition for Next Generation Life Science.
- Database and/or data collection tool is easy to use and readily available to all team members.
-

### Project Constraints

#### Time

- Career outcome collection and classification must be completed by 9/15/2017, as Student Inten/Project Support is only available from 7/17/2017 to 9/15/2017.
- Project team members have limited weekly time to devote to data collection, consolidation and analysis.
  - Project Manager- 40 hours per week
  - Project Support- 32 hours per week
  - SME(s) – 5-10 hours per week

#### Scope

- Data collection and analysis is limited to UCSF graduate students. A separate project to collect and analyze postdoctoral career outcomes may be initiated, depending upon the success and results of this project.

#### Budget

- Cost of project management and support resources should not exceed original estimation
- There is an incentive to maintain a relatively low total project cost, to demonstrate the minimal expense it will take to collect, analyze and make available to the UCSF community.

Created by UCSF Graduate Division

### Project Mangaement Approach

#### Project Organization

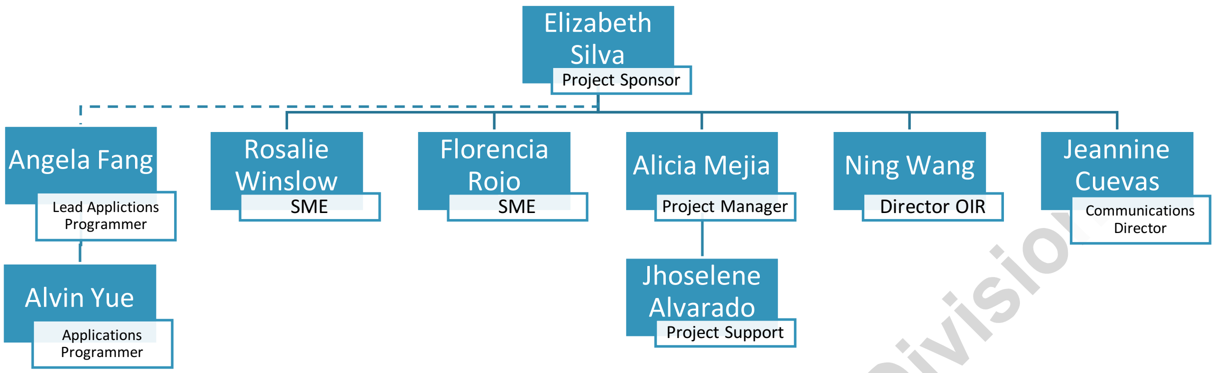

| Role | High-level Responsibilities |
| --- | --- |
| **Project Sponsor** | • Decision maker for overall project<br>• Enable project director and manager in collecting and analyzing data<br>• Participate as a data auditor |
| **SME** | • Decision maker for what type of data to collect and how to analyze it<br>• Key point of contact for programs and administrators<br>• Participate as a data auditor<br>• Secondary role to collect and consolidate data |
| **Project Manager** | • Project management- document project goals, document and communicate project status, track time and effort spent, roles & responsibilities, and other project monitoring duties.<br>• Data collection, consolidation and management in REDCap as secondary role.<br>• Database administration and data quality audits/cleanup.<br>• Internal and external Organizational Change Management |
| **Project Support** | • Primary role to collect and consolidate career outcomes in REDCap |
| **Applications Programmer(s)** | • Data extraction, collection and reporting of trainee program/appointment demographic information from registrar, for API upload into the REDCap database.<br>• Creation and maintenance of REDCap API import of quarterly updates of trainee information. |
| **Director of Institutional Research** | • Analysis of trainee information and career outcomes for creation and publication into Tableau dashboard |
| **Communications Director** | • Coordination with Director of Institutional Research to ensure Tableau presentation is within UCSF brand identity guidelines. |

Created by UCSF Graduate Division

| Role | High-level Responsibilities |
| --- | --- |
| | • Website development to publish Tableau dashboards and tabular data on appropriate websites. |

#### RACI

Greater detail regarding project roles and responsibilities of each team member and stakeholder group by phase of the project will be documented and communicated to the project team.

#### Governance and Decision-Making Procedures/Norms

- Project team members meet on a regular interval to discuss timeline, deliverables, issues, decisions, risks and other topics related to the completion of the project.
- Project Sponsor is the overall responsible and ultimate decision maker for the project, delegation of deliverables, and budget.
- The Project Sponsor and SME(s) make decisions about what data to collect and how to analyze data. Project Sponsor and SME(s) are also the key points of contact for programs and administrators.
- The Project Manager will complete follow-up activities to ensure team is on-task, and provide a weekly project status report to the Project Sponsor or other relevant stakeholders.

#### Project Communication Plan

The Communication Plan for project communication and collaboration both internal and external to the project team will be documented and approved by the project team members.

#### Project Monitor

Project Risks, Assumptions, Issues, Dependencies (RAID), Action Items, and Decisions will be documented and communicated by the Project Manager as the project progresses. Details on these logs are located in the Project Communication Plan.

- *Action Log*
- *Decision Log*
- *RAID:*
  - *Risk Log*
  - *Assumptions*
  - *Issue Log*
  - *Dependencies*

Created by UCSF Graduate Division

#### Organizational Change Management Approach

The purpose of this project is to pilot a more robust collection and analysis of graduate student career outcomes. It proposes a change from the current state to a desired future state in which more reliable career trajectory information is collected and shared with graduate students, the UCSF community, other institutions and the general public. The goal is to influence other academic institutions to collect and share their student career outcomes information. An Organizational Change Management plan will be drafted to include stakeholder analysis, external communication plan, training, and change execution risk management. This will ensure that we undergo the necessary activities to obtain buy-in from impacted stakeholders, accelerate the adoption of these processes, and sustain these new behaviors beyond the closure of this project.

### Institutionalization of Ongoing Career Outcome Collection

#### Sustainment Strategy

##### 1. Sustainment Deliverables-

- Outline specific deliverables required to maintain ongoing career outcome collection, analysis and publication, as well as database management/administration.

##### 2. Sustainment Cost-

- Determine staffing, consulting and/or other resource costs required for the sustainment of the database(s) and career outcome collection and presentation.

##### 3. Roles and Responsibilities-

- Key roles and responsibilities for the development and execution of the ongoing solution.

##### 4. User Groups-

- List of end-users for your database(s), with a high-level summary of the type of support required and number of impacted users in each group.

##### 5. Training-

- Determine training and support required for user groups and other career outcomes stakeholders.

##### 6. Support Services-

- A high-level description of inter-departmental and/or consulting support required to maintain database and career outcome collection and communication, including relevant time frames.

##### 7. Communication Plan-

- A robust communication plan to designate essential intra- and inter-departmental and/or organizational communications, included designated audience, frequency, source/medium, accountable/responsible team member(s), and relevant approvals.

Created by UCSF Graduate Division
