## Supplemental file 6 for "Where do our graduates go? A toolkit for retrospective and ongoing career outcomes data collection for biomedical PhD students and postdoctoral scholars"

### Career outcomes job categories and definitions

| Sector | |
|---|---|
| Academia | Academic institutions of higher education, including colleges, universities, some medical centers, or free-standing research institutions where training occurs. This does not include VA hospitals, but does include teaching, for-profit, and other types of hospitals. |
| Government | Any organization operated by federal, state, local or foreign governments. Includes VA hospitals. |
| For-Profit | Any organization that operates to make a profit, including some industry research. |
| Nonprofit | Any non-governmental organization that does not operate to make a profit. Includes K-12 institutions. |
| Other | Individuals who are unemployed, full-time caretaker or parent, on extended medical leave or employed at an organization not included in other options. |
| Unknown | Unknown |
| N/A | Not applicable |

| Career Types | |
|---|---|
| Primarily Research | The primary, although not necessarily the only, focus is the conduct or oversight of scientific research. Includes academic faculty titles at R1-R3 institutions as identified through Carnegie classifications. |
| Primarily Teaching | The primary, although not necessarily the only, focus is education and teaching. Includes academic faculty at all other institutions. |
| Science-related | Career that is relevant to the conduct of scientific research, but does not directly conduct or oversee research activities. |
| Not Related to Science | Career that is not directly relevant to the conduct of scientific research. |
| Further Training or Education | Temporary training position. Examples include: postdoctoral research, completing medical residency, or pursuing an additional degree. |
| Unknown | Unknown |
| N/A | Not applicable |

| Job Function | |
|---|---|
| Administration | Administrative-intensive roles. Examples include: Faculty affairs, graduate program administrators, human resources, academic admissions, career development offices, grant and contracts management, research development, PhD-level program development. |
| Business Development, Consulting, and Strategic Alliances | Role that involves the development, execution, management, or analysis of a business. Role may include relationship management, refinement of operational efficiency, or fee-based advisory services. Examples include management consultant, business development professional, market researcher, investment analyst, venture capitalist. |

| Job Function | |
|---|---|
| Clinical Research Management | Role that is responsible for the oversight, management, or design of clinical research trials. Examples include clinical research project/trials manager or coordinator. |
| Clinical Services | Role that involves that administration of clinical services or research. Examples include genetics counselor, testing specialist, and clinical laboratory staff. |
| Data Science, Analytics, and Software Engineering | Role that may combine programming, analytics, advanced statistics, data communication, and/or software development. |
| Entrepreneurship | Founder, co-founder, CEO or other role that develops, manages, and provides/obtains capital to initiate a business or enterprise. This function does not include staff at a start-up business. |
| Faculty: Nontenure Track | Leading an academic research team and ineligible for tenure. Examples include: Research assistant professor, research associate professor, research professor. |
| Faculty: Tenured/Tenure Track | Leading an academic research team and eligible for or already tenured. Examples include: assistant professor, associate professor, professor. |
| Faculty: Tenure Track Unclear or Not Applicable | Leading an academic research team at an institution where tenure is not granted or tenure status is unknown. For those tracking down alumni and binning them into job functions, whether someone is or is not on a tenure track is often not clear and should be sorted here. |
| Full-time Teaching Staff (Instructor/Lecturer) | Full-time institutionalized teaching position with no research responsibilities. Examples include Instructor, Lecturer. Distinct from "Primarily teaching, faculty," these are people teaching at a single university without a faculty appointment. |
| Group Leader (Research) | Leading a research team in a nonacademic setting. Anyone working in industry, non-profit or government who is running a somewhat independent research group. This includes those with "Faculty" titles at VA hospitals and other government research institutions. |
| Healthcare Provider | Role where the primary responsibility is providing healthcare. Examples include doctor, nurse, medical residents, and veterinarian. |
| Intellectual Property and Law | Role that involves the curation, management, implementation or protection of intelligence and creation, including trademarks, copyrights, patents, or trade secrets. Examples include patent agent, patent attorney, and technology transfer specialist. |
| Part-time Teaching Staff (Adjunct) | Contingent teaching role that is contracted on a single-semester, short-term, or non-permanent basis with no research responsibilities. Examples include instructor, lecturer. Distinct from "Primarily teaching, faculty," these could include people teaching at multiple universities, indicating contingent status. |
| Postdoctoral (Research) | Temporary mentored training position in scientific research environment following completion of doctoral degree. |
| Regulatory Affairs | Role that involves controlling or evaluating the safety and efficacy of products in areas including pharmaceuticals, medicines, and devices. Examples include institutional regulatory affairs professional, quality control specialist, compliance officer. |

| Job Function | |
|---|---|
| Research Staff or Technical Director | Role that directly involves performing or managing research. Examples include research staff, staff scientists, lab/core managers, directors of research facilities, public health analyst, and epidemiologists. |
| Sales and Marketing | Non-technical role that is related to the sales or marketing of a science-related product or service. Examples include medical science liaison, technical sales representative, and marketing specialist. |
| Science Education and Outreach | Role that involves K-12 teaching or public outreach at a primary/secondary schools, science museum, scientific society, or similar. Examples include high school teacher, museum curriculum development, outreach program administrator. |
| Science Policy and Government Affairs | Role that involves policy or program development and review, including analysis, advisory, or advocacy. Examples include program officer, public affairs or government affairs staff at scientific societies, foundations, government entities, or think tanks. |
| Science Writing and Communication | Role that involves the communication of science-related topics. Examples include science, medical, or technical writer, journalist, science editor, science publisher. |
| Technical Support and Product Development | Role that requires specialized technical knowledge of a science-related product. Examples include technical support specialist, field application specialist, product development scientist or engineer. |
| Other | Role that does not require scientific training or involve the direct implementation or communication of science. Examples include full-time homemaker, caretaker, chef, food or hospitality services, some types of military service or mission work, or currently unemployed. |
| Completing Further Education | Pursuing additional education that usually results in graduation with conferment of a degree or certificate; this does *not* include postdoctoral research. Examples include: pursuing an additional degree in medicine, law, business, or other area. |
| Deceased/retired | Deceased or retired |
| Unknown | Unknown |
| N/A | Not applicable |

### Career Types – Guidance Grids

| Legend: |
|---|
| Appropriate for Career Type |
| Not appropriate for Career Type |
| Further training or education non-Postdoctoral |

| Academia | Primarily research | Primarily teaching | Science-related | Further training or education | Not related to science | Unknown |
|---|---|---|---|---|---|---|
| Administration | Not appropriate | Not appropriate | Appropriate | Further training | Not appropriate | Not appropriate |
| Business Development, Consulting, and Strategic Alliances | Not appropriate | Not appropriate | Appropriate | Further training | Not appropriate | Not appropriate |
| Clinical Research Management | Not appropriate | Not appropriate | Appropriate | Further training | Not appropriate | Not appropriate |
| Clinical Services | Not appropriate | Not appropriate | Appropriate | Further training | Not appropriate | Not appropriate |
| Data Science, Analytics, and Software Engineering | Appropriate | Not appropriate | Appropriate | Further training | Not appropriate | Not appropriate |
| Entrepreneurship | Not appropriate | Not appropriate | Appropriate | Further training | Not appropriate | Not appropriate |
| Faculty member - Nontenure track | Appropriate | Appropriate | Not appropriate | Not appropriate | Not appropriate | Not appropriate |
| Faculty member - Tenure/Tenure track | Appropriate | Appropriate | Not appropriate | Not appropriate | Not appropriate | Not appropriate |
| Faculty member – Track unclear or not applicable | Appropriate | Appropriate | Not appropriate | Not appropriate | Not appropriate | Not appropriate |
| Full-time teaching staff (Instructor/Lecturer) | Not appropriate | Appropriate | Not appropriate | Further training | Not appropriate | Not appropriate |
| Healthcare Provider | Not appropriate | Not appropriate | Appropriate | Appropriate | Not appropriate | Not appropriate |
| Intellectual Property and Law | Not appropriate | Not appropriate | Appropriate | Further training | Not appropriate | Not appropriate |
| Part-time teaching staff (Adjunct) | Not appropriate | Appropriate | Not appropriate | Further training | Not appropriate | Not appropriate |
| Postdoctoral (research) | Not appropriate | Not appropriate | Not appropriate | Appropriate | Not appropriate | Not appropriate |
| Regulatory Affairs | Not appropriate | Not appropriate | Appropriate | Further training | Not appropriate | Not appropriate |
| Research Staff or Technical Director | Appropriate | Not appropriate | Not appropriate | Not appropriate | Not appropriate | Not appropriate |
| Sales and Marketing | Not appropriate | Not appropriate | Appropriate | Further training | Not appropriate | Not appropriate |
| Science Education and Outreach | Not appropriate | Not appropriate | Appropriate | Further training | Not appropriate | Not appropriate |
| Science Policy and Government Affairs | Not appropriate | Not appropriate | Appropriate | Further training | Not appropriate | Not appropriate |
| Science Writing and Communication | Not appropriate | Not appropriate | Appropriate | Further training | Not appropriate | Not appropriate |
| Technical Support and Product Development | Not appropriate | Not appropriate | Appropriate | Further training | Not appropriate | Not appropriate |
| Completing further education | Not appropriate | Not appropriate | Not appropriate | Appropriate | Not appropriate | Not appropriate |
| Other | Not appropriate | Not appropriate | Not appropriate | Not appropriate | Appropriate | Not appropriate |
| Unknown | Appropriate | Appropriate | Appropriate | Appropriate | Appropriate | Appropriate |

| Government | Primarily research | Primarily teaching | Science-related | Further training or education | Not related to science | Unknown |
|---|---|---|---|---|---|---|
| Administration | Not appropriate | Not appropriate | Appropriate | Further training | Not appropriate | Not appropriate |
| Business Development, Consulting, and Strategic Alliances | Not appropriate | Not appropriate | Appropriate | Further training | Not appropriate | Not appropriate |

| **Government** | Primarily research | Primarily teaching | Science-related | Further training or education | Not related to science | Unknown |
|---|---|---|---|---|---|---|
| Clinical Research Management | | | | | | |
| Clinical Services | | | | | | |
| Data Science, Analytics, and Software Engineering | | | | | | |
| Entrepreneurship | | | | | | |
| Group Leader (research) | | | | | | |
| Healthcare Provider | | | | | | |
| Intellectual Property and Law | | | | | | |
| Postdoctoral (research) | | | | | | |
| Regulatory Affairs | | | | | | |
| Research Staff or Technical Director | | | | | | |
| Sales and Marketing | | | | | | |
| Science Education and Outreach | | | | | | |
| Science Policy and Government Affairs | | | | | | |
| Science Writing and Communication | | | | | | |
| Technical Support and Product Development | | | | | | |
| Completing further education | | | | | | |
| Other | | | | | | |
| Unknown | | | | | | |

| **For-profit** | Primarily research | Primarily teaching | Science-related | Further training or education | Not related to science | Unknown |
|---|---|---|---|---|---|---|
| Administration | | | | | | |
| Business Development, Consulting, and Strategic Alliances | | | | | | |
| Clinical Research Management | | | | | | |
| Clinical Services | | | | | | |
| Data Science, Analytics, and Software Engineering | | | | | | |
| Entrepreneurship | | | | | | |
| Group Leader (research) | | | | | | |
| Healthcare Provider | | | | | | |
| Intellectual Property and Law | | | | | | |
| Postdoctoral (research) | | | | | | |
| Regulatory Affairs | | | | | | |
| Research Staff or Technical Director | | | | | | |
| Sales and Marketing | | | | | | |
| Science Education and Outreach | | | | | | |

| **For-profit** | Primarily research | Primarily teaching | Science-related | Further training or education | Not related to science | Unknown |
|---|---|---|---|---|---|---|
| Science Policy and Government Affairs | | | | | | |
| Science Writing and Communication | | | | | | |
| Technical Support and Product Development | | | | | | |
| Completing further education | | | | | | |
| Other | | | | | | |
| Unknown | | | | | | |

| **Nonprofit** | Primarily research | Primarily teaching | Science-related | Further training or education | Not related to science | Unknown |
|---|---|---|---|---|---|---|
| Administration | | | | | | |
| Business Development, Consulting, and Strategic Alliances | | | | | | |
| Clinical Research Management | | | | | | |
| Clinical Services | | | | | | |
| Data Science, Analytics, and Software Engineering | | | | | | |
| Entrepreneurship | | | | | | |
| Group Leader (research) | | | | | | |
| Healthcare Provider | | | | | | |
| Intellectual Property and Law | | | | | | |
| Postdoctoral (research) | | | | | | |
| Regulatory Affairs | | | | | | |
| Research Staff or Technical Director | | | | | | |
| Sales and Marketing | | | | | | |
| Science Education and Outreach | | | | | | |
| Science Policy and Government Affairs | | | | | | |
| Science Writing and Communication | | | | | | |
| Technical Support and Product Development | | | | | | |
| Completing further education | | | | | | |
| Other | | | | | | |
| Unknown | | | | | | |

| **Other** | Primarily research | Primarily teaching | Science-related | Further training or education | Not related to science | Unknown |
|---|---|---|---|---|---|---|
| Administration | | | | | | |
| Business Development, Consulting, and Strategic Alliances | | | | | | |
| Clinical Research Management | | | | | | |
| Clinical Services | | | | | | |

| **Other** | Primarily research | Primarily teaching | Science-related | Further training or education | Not related to science | Unknown |
|---|---|---|---|---|---|---|
| Data Science, Analytics, and Software Engineering | | | | | | |
| Entrepreneurship | | | | | | |
| Group Leader (research) | | | | | | |
| Healthcare Provider | | | | | | |
| Intellectual Property and Law | | | | | | |
| Postdoctoral (research) | | | | | | |
| Regulatory Affairs | | | | | | |
| Research Staff or Technical Director | | | | | | |
| Sales and Marketing | | | | | | |
| Science Education and Outreach | | | | | | |
| Science Policy and Government Affairs | | | | | | |
| Science Writing and Communication | | | | | | |
| Technical Support and Product Development | | | | | | |
| Completing further education | | | | | | |
| Other | | | | | | |
| Unknown | | | | | | |
